# Reduced competence to arboviruses following the sustainable invasion of *Wolbachia* into native *Aedes aegypti* from Niterói, Southeastern Brazil

**DOI:** 10.1101/2020.09.25.312207

**Authors:** João Silveira Moledo Gesto, Gabriel Sylvestre Ribeiro, Marcele Neves Rocha, Fernando Braga Stehling Dias, Julia Peixoto, Fabiano Duarte Carvalho, Thiago Nunes Pereira, Luciano Andrade Moreira

## Abstract

Field release of *Wolbachia*-infected *Aedes aegypti* has emerged as a promising solution to manage the transmission of dengue, Zika and chikungunya in endemic areas across the globe. Through an efficient self-dispersing mechanism, and the ability to induce virus-blocking properties, *Wolbachia* offers an unmatched potential to gradually modify wild *Ae. aegypti* populations turning them unsuitable disease vectors. Here in this work, a proof-of-concept field trial was carried out in a small community of Niterói, greater Rio de Janeiro. Following the release of *Wolbachia*-infected eggs, we reported a successful invasion and long-term establishment of the bacterium across the territory, as denoted by stable high-infection indexes (>80%). We have also demonstrated that refractoriness to dengue and Zika viruses, either thorough oral-feeding or intra-thoracic saliva challenging assays, were maintained over the adaptation to the natural environment of Southeastern Brazil. These findings further support *Wolbachia*’s ability to invade local *Ae. aegypti* populations and impair disease transmission, and shall pave the way for future epidemiological and economic impact assessments.

## Introduction

The mosquito *Aedes aegypti* (= *Stegomyia aegypti*) holds a core status among tropical disease vectors, being able to host and transmit a broad variety of viruses, such as those causing dengue, Zika and chikungunya (Kraemer et al., 2015; WHO, 2017). Dengue virus (DENV) is certainly the most prevalent, with a global distribution spectrum including 128 countries (Brady et al., 2012) and approximately 400 million infections annually (Bhatt et al., 2013). Brazil accounts for most of these cases, with more than 1.5 million infections only in 2019 (SVS, 2019). In spite of being historically less prevalent, chikungunya (CHIKV) and Zika (ZIKV) viruses, restricted to Africa and Asia until the early 2000’s, were recently emerging in new territories and augmenting their impact on public health (Mayer et al., 2017; Nugent et al., 2017; Wahid et al., 2017; Weaver, 2014). The introduction of CHIKV in Central and South America, for example, led to about one million suspected disease cases between 2013 and 2014 (McSweegan et al., 2015). Similarly, ZIKV first records in America date to the end of 2013, after which it rapidly spread through the continent (Faria et al., 2016). In 2015, the World Health Organization (WHO) declared a Public Health Emergency of International Concern following a serious ZIKV outbreak in Northeast Brazil, associated with high rates of microcephaly in newborns (Barreto et al., 2016).

Since effective vaccines or therapeutic drugs are still under development (Abdelnabi et al., 2015; Lin et al., 2018; Thisyakorn and Thisyakorn, 2014) and not currently available to fight outbreaks of Zika, dengue and chikungunya, public health initiatives rely entirely on vector control. Traditionally, this is achieved by the mechanical elimination of breeding sites and the use of chemical insecticides to reduce *Ae. aegypti* populations. However, both methods have proven inefficient and unsustainable for the long term, mostly due to the multitude of artificial breeding sites used by this species in urban landscapes (Carvalho and Moreira, 2017; Valença et al., 2013) and the advent of naturally resistant allelic variants (Linss et al., 2014; Maciel-de-Freitas et al., 2014; Moyes et al., 2017). In face of these challenges, the development of new solutions is, therefore, a critical need for a more efficient control of *Ae. aegypti* populations and/or disease transmission.

In recent years, an innovative approach using the endosymbiont *Wolbachia pipientis* to block arbovirus transmission has been proposed and successfully tested in *Ae. aegypti* (Dutra et al., 2015; Hoffmann et al., 2014, 2011; Walker et al., 2011), gaining momentum as a viable and sustainable alternative to traditional control methods. Naturally found in around 40% of all arthropods (Zug and Hammerstein, 2012), *Wolbachia* are maternally-inherited intracellular bacteria which exploits host reproductive biology to increase its transmission rates and dispersal in nature. For some bacterial strains, this is achieved by triggering a phenomenon called cytoplasmic incompatibility (CI), which turns the progeny unviable when an infected male copulates with an uninfected female (Werren et al., 2008). Remarkably, *Wolbachia*-host association also leads to pathogen interference (PI) phenotypes (Lindsey et al., 2018; Teixeira et al., 2008), particularly relevant to vector control applied studies. In view of this, stable and heritable lines, harboring different *Wolbachia* strains, have been created following artificial transinfection of *Ae. aegypti* (Ant et al., 2018; McMeniman et al., 2009; McMeniman and O’Neill, 2010; Moreira et al., 2009; Walker et al., 2011). Through pathways involving the modulation of host immune system (Rancès et al., 2012) and metabolite sequestration (Caragata et al., 2014; Geoghegan et al., 2017), *Wolbachia*-harboring lines exhibit high-levels of refractoriness to DENV, CHIKV, ZIKV, and other medically relevant arboviruses (Aliota et al., 2016a, 2016b; Dutra et al., 2016; Moreira et al., 2009). As such, releasing some of these lines in the field to gradually replace virus-susceptible wild populations could potentially reduce transmission and human infection rates.

Successful field release trials using *Ae. aegypti* lines infected with *Wolbachia* wMel strain have been reported in Northern Australia, Vietnam and more recently in Southeastern Brazil (Garcia et al., 2019; Hoffmann et al., 2011). In the latter, a small community in Rio de Janeiro (RJ) was subject to a rolling out strategy based on adult releases, which led to the invasion and long-term establishment of the bacterium in the field. Nonetheless, important vector competence data following the invasion is still lacking and needs to be assessed to in order to provide evidence of virus-blocking maintenance.

In this study, we report the results of a field release trial in Jurujuba, a small suburb of Niterói (Rio de Janeiro State), in which eggs were used for introducing *Wolbachia* into *Ae. aegypti* wild specimens. Besides evaluating *Wolbachia*’s ability to invade mosquito populations, we investigated the bacterium density level and the vector competence for ZIKV and DENV in post-release field samples, contributing to a better characterization of targeted populations in Southern Brazil.

## Results

### Egg releases successfully promote *Wolbachia* invasion in Jurujuba, Niterói (RJ)

To evaluate the efficiency of egg releases as a method of *Wolbachia* field deployment in Brazil, we carried out a pilot study in Jurujuba, a suburban area of Niterói (Rio de Janeiro State). Using Mosquito Release Containers (MRCs), *Wolbachia*-infected eggs (*w*MelRio strain) were mixed with standard amounts of food and water, and distributed over all seven Jurujuba’s sectors in a timely fashion (Figure 1, Supplementary Figure 1, Supplementary Table 1). BG-sentinels were also spatially distributed (Supplementary Figure 2, Supplementary Table 2) and used to collect *Ae. aegypti* field specimens for *Wolbachia* molecular diagnosis, and calculation of infection rates (percent infected individuals).

**Figure 1.**
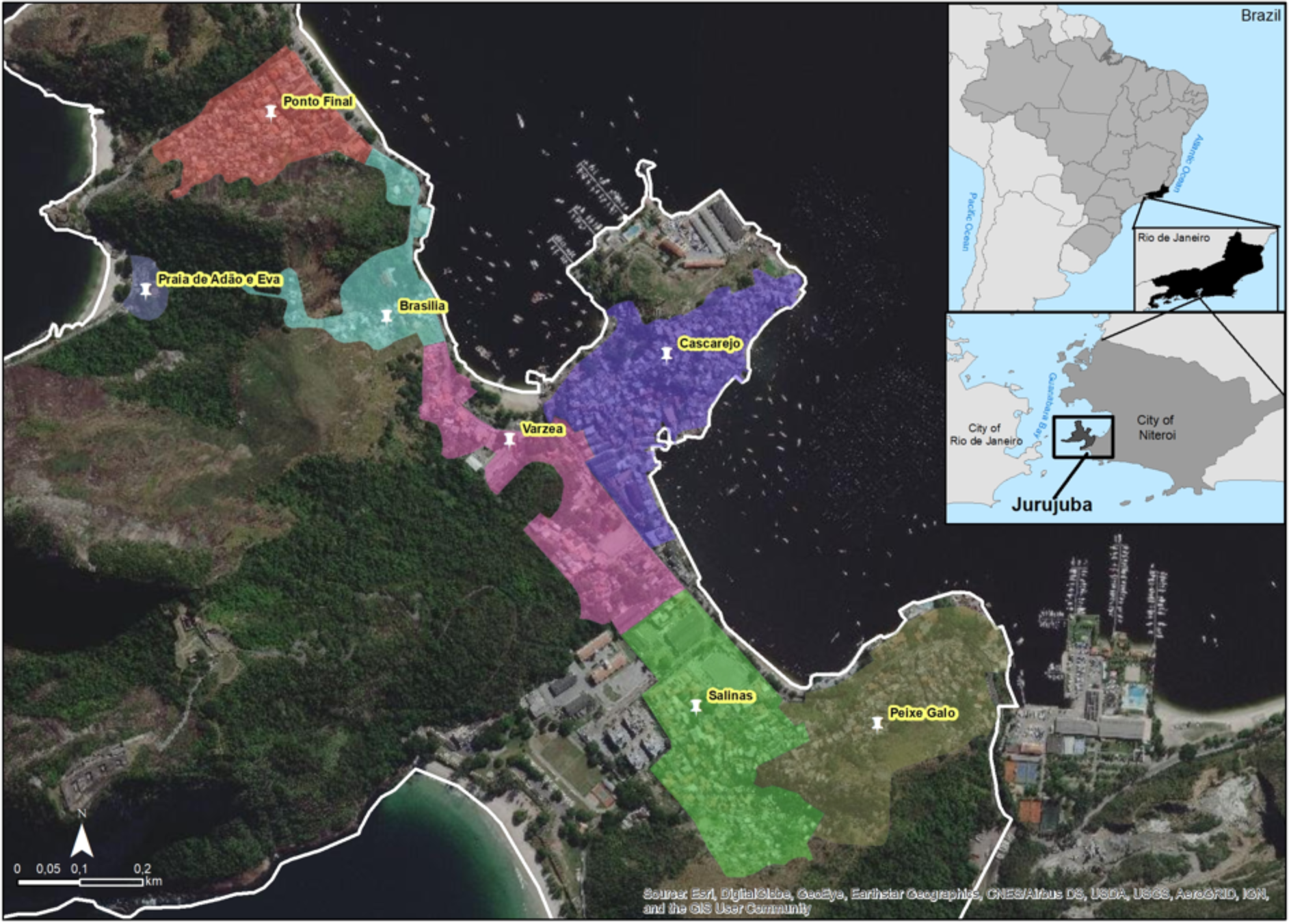
Map of Jurujuba release areas. Satellite view of Jurujuba, a suburban neighborhood of Niterói (RJ). With an estimated population of 2,797 in 2.53 km^2^, Jurujuba was divided into seven release areas (highlighted) according to local sectors: Ponto Final, Várzea, Brasília, Cascarejo, Praia de Adão e Eva, Peixe-Galo and Salinas.

Situated at the furthermost housing area of Jurujuba’s peninsula, Ponto Final was selected the starting sector for field release, from which adjacent ones would derive following a rollout strategy. As such, all related activity planning, including community engagement, territorial mapping, release and monitoring sites allocation, and mass-rearing production scale was initially tuned to Ponto Final. Here, *Wolbachia*-infected eggs were released over 25 consecutive weeks, from August 2015 to February 2016, during which infection rates rapidly increased from basal to high levels (>80%) (Figure 2). In fact, by week 17, infection rates hit 81% and, by the end of the release period, 88%. Post-release infection rates were maintained at high levels in subsequent months, suggesting a successful invasion at this particular sector and paving the way to cover others, fulfilling our initial design.

**Figure 2.**
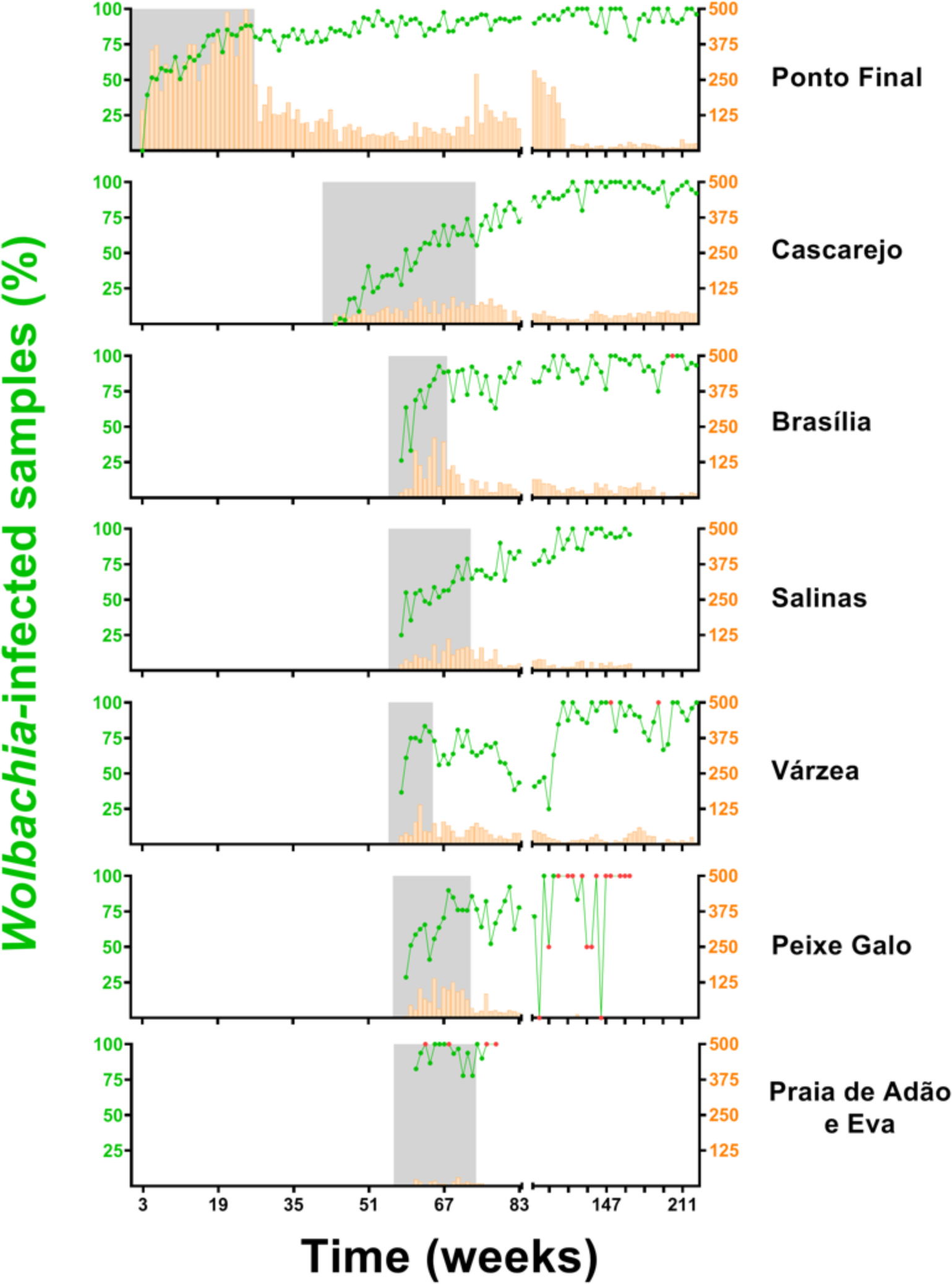
*Wolbachia*’s spread across Niteroi’s suburb Jurujuba. Over a period spanning 73 weeks, *w*Mel-infected eggs were gradually released across all sectors of Jurujuba: Ponto Final, Cascarejo, Brasília, Salinas, Várzea, Peixe Galo and Praia de Adão e Eva (release interventions depicted under grey shaded areas). Infection indexes (%) are plotted as line-connected dots (green), following the vertical left axis, whereas the sample sizes are plotted as histograms (orange), following the vertical right axis. Small-sized samples (N<5) were marked in red. The horizontal axis represents time, in number of weeks, since the beginning of egg-releases in Ponto final (August 2015) until more recent days (December 2019). Note that the horizontal axis is split into two fragments, in which the data accounts for 1-week (left fragment) or 4-week bins (right fragment). Ticks are scaled accordingly to represent a 16-week bin.

Following new planning and schedules, the release of *Wolbachia*-infected eggs was extended to Cascarejo, in June 2016, and to Brasília, Salinas, Várzea, Peixe-Galo and Praia de Adão e Eva, in September 2016 (Figure 2). The release period duration and number of MRCs allocated in each sector varied slightly (Supplementary Table 1), due to their intrinsic properties (i.e. housing area and human population density). By mid-January 2017, egg release ceased in whole Jurujuba, with most sectors recording mid-to-high rates of *Wolbachia* infection (60-90%). Mimicking Ponto Final, the post-release phase generally followed the positive trends of the release period and featured increasing infection rates towards near fixation levels (90-100%) (Figure 2). This was clearly the case for Cascarejo, Brasília and Salinas. A notable exception was Várzea, which recorded an erratic invasion profile, with high infection rates by the end of the release period (i.e. ∼ 80% in 8 weeks) and a gradual decrease over the following months, when rates hit less than 40%. Here, a complete recovery and stability at high rates was achieved only after 10 months into the post-release phase. Peixe Galo also revealed a slightly erratic profile but, unlike Várzea, it did not resemble a negative trend. Instead, mid-to-high level oscillation was sustained over the post-release phase, and part of the noise might be attributed to low sampling. Lastly, Praia de Adão e Eva exhibited consistently high rates over the release period and the following few weeks, suggesting a standard invasion profile. However, given its relatively small area and population, post-release monitoring was suspended, yielding no long-term data and preventing a more assertive analysis for this sector.

By aggregating weekly infection rates from different sectors, an overall post-release profile of Jurujuba was revealed (Figure 3). Confirming previous data analysis, in which sectors were individually treated, the overall profile revealed that Jurujuba consistently recorded high infection rates (80-100%) along the post-release phase, from mid-January 2017 until December 2019. Data encompassing this period, of almost three years, suggest that *Wolbachia* successfully invaded Jurujuba, being able to sustain infection rates in the long-term.

**Figure 3.**
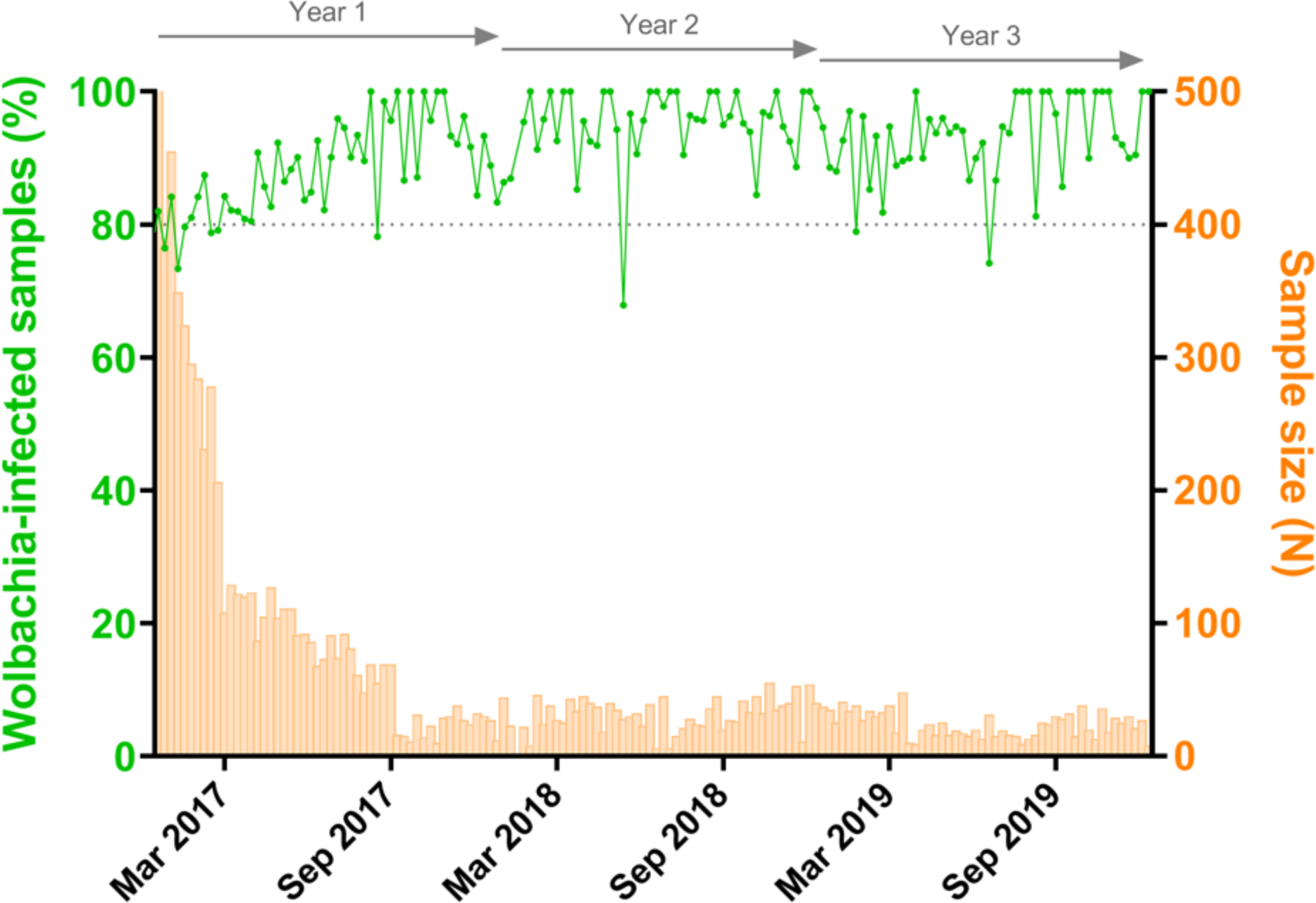
Post-release establishment of *Wolbachia* in Jurujuba. Following egg-release cessation in Jurujuba, *Wolbachia-*infection was monitored over the following three years, during which a self-sustained field establishment was observed. Infection indexes (%) are observed as line-connected dots (green), following the vertical left axis, whereas the sample sizes are plotted as histograms (orange), following the vertical right axis. The horizontal axis denotes time (weeks), with ticks scaled for 6-month bins. Top arrows mark yearly post-release periods. The dashed line, positioned at the 80% index mark, denotes a threshold for high-percent infection.

### *Wolbachia* density is higher in post-release field samples

To investigate whether *Wolbachia* whole-body density (i.e. titer) was affected over time in Jurujuba’s environment, field samples were tested a few months after release cessation (March-May 2017) and one year later (March-May 2018). Counterpart *w*MelRio colony samples (i.e. collected at equivalent time periods) were also tested to assess lab-rearing data and control generation-dependent variation in density, which could possibly mask evolutionary changes in this trait. Density data were plotted side-by-side to reveal putative differences between field and colony samples, as well as details of their distribution (Figure 4). Indeed, visual inspection suggests that *Wolbachia* density vary among groups, which was further corroborated by statistical analysis (*H* = 340.2, *P* < 0.0001). Subsequent multiple comparisons revealed that ‘field’ densities are higher than ‘colony’ ones, in samples both from the beginning (*P* < 0.0001) or from ∼ 1 year into the post-release phase (*P* < 0.0001). Moreover, when field samples are compared to themselves, a significant increase in density over time is detected (*P* < 0.0001), suggesting an evolving *Wolbachia*-host relationship in Jurujuba’s environment.

**Figure 4.**
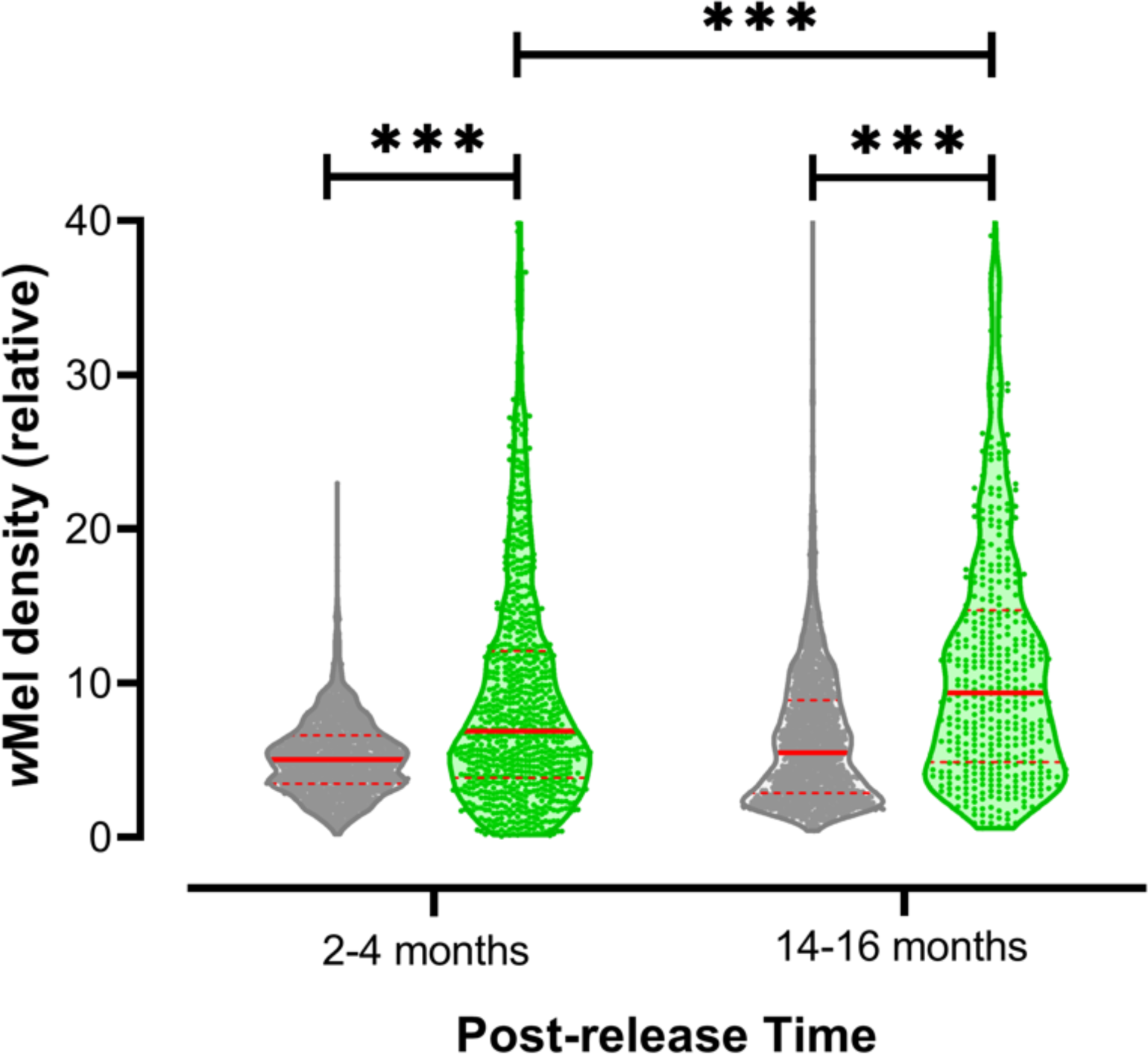
*Wolbachia* whole-body density increases subsequent to field establishment. *Wolbachia* whole-body density was molecularly assayed in whole-body extracts from colony (grey) and Jurujuba samples (green), collected over the post-release phase. Density or titration levels (vertical axis) are relative quantification indexes, reflecting the copy number ratio between *w*Mel WD0513 and the endogenous ribosomal maker RPS17. Data were aggregated in 12-weeks pools and represented by violin plots, with medians and quartiles in solid and dashed red lines, respectively. Statistical inferences were performed by the non-parametric Kruskal-Wallis test followed by Dunn’s post-hoc multiple comparisons. Asterisks highlight differences between groups, considering a significance level (α) equal to 0.05, and its numbers reflects probability ranges: *P* < 0.05 = *, *P* < 0.01 = **, *P* < 0.001 = ***.

### *Wolbachia* inhibits DENV e ZIKV replication in the head and thorax of field samples

Following the invasion and long-term stability of *Wolbachia* in Jurujuba, *Ae. aegypti* field samples were submitted to vector competence assays. Jurujuba specimen eggs were collected in Ponto Final over three months, from April to June 2017, which correspond to 14-16 months into the post-release phase of this particular sector. Specimens eggs from Urca, a *Wolbachia*-free area in the neighboring city, Rio de Janeiro, were collected at the same time period and served as experimental controls. F1 adult females, from Jurujuba and Urca, were orally challenged with ZIKV or DENV, and viral titers were assessed 14 days post infection (dpi) in head/thorax individual extracts. Two independent assays were performed for each virus.

Our results revealed that oral challenging with ZIKV could trigger infection and high viral titers in most samples from Urca, but could not elicit a similar outcome in samples from Jurujuba, which mostly failed to infect (Figure 5A). Statistical analyses corroborate the significant reduction of ZIKV titers in Jurujuba samples, in both first (*U* = 47.5, *P* < 0.0001) and second assays (*U* = 36.5, *P* < 0.0001). The oral challenging with DENV led to an almost identical picture, with most samples from Urca being infected with high viral titers, whilst in samples from Jurujuba only a few were infected (Figure 5B). Once again, significant differences between Urca and Jurujuba were found in the first (*U* = 74, *P* < 0.0001) and second assays (*U* = 20.5, *P* < 0.0001). In addition, we double-checked the *Wolbachia* status of the same F1 samples, further confirming its majoritarian presence in Jurujuba (∼ 88% rate; 55 out of 62 individuals tested positive for *Wolbachia*) and complete absence in Urca (Supplementary Figure 3). Altogether, our data suggest that the oral exposure with ZIKV or DENV is less prone to trigger and disseminate infection in *Wolbachia*-harboring Jurujuba specimens than in *Wolbachia*-negative Urca correlates, and that this effect is bacterium-driven.

**Figure 5.**
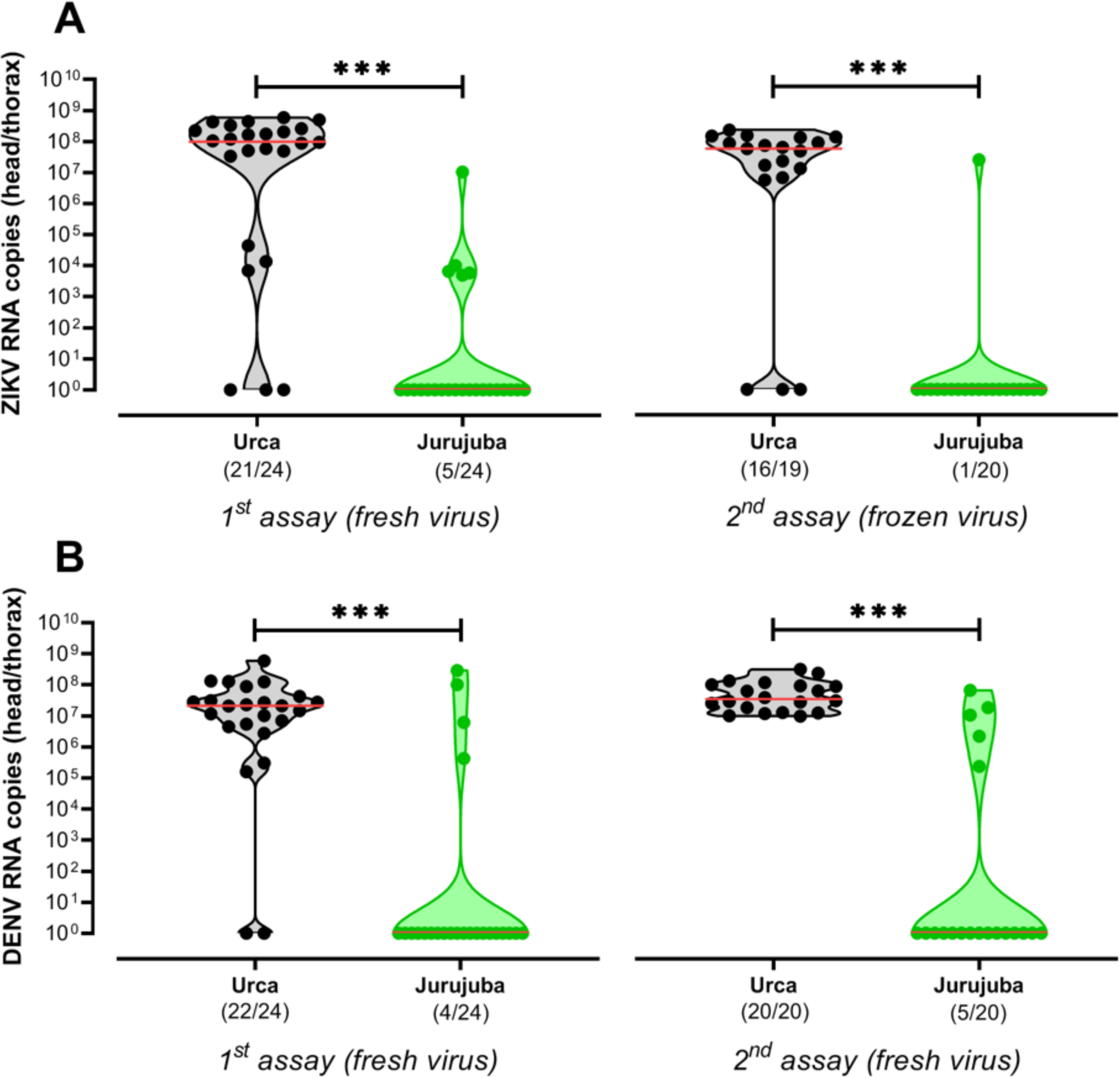
*Wolbachia* inhibits ZIKV and DENV tissue replication following oral-infection of field samples. Following the long-term establishment of *Wolbachia* in Jurujuba, field samples were brough to the laboratory and orally challenged with ZIKV and DENV, either fresh or frozen. *Wolbachia*-mediated inhibition of ZIKV and DENV replication was evaluated by comparing samples from Jurujuba (green) and Urca (grey), a *Wolbachia*-free non-targeted area in the suburbs of Rio de Janeiro. Violin plots represent absolute quantifications of (A) DENV and (B) ZIKV titers in head/thorax extracts, at 14 dpi. Median and quartiles are depicted in solid and dashed red lines, respectively. Non-parametric Mann-Whitney U test highlighted differences between Urca and Jurujuba. Asterisks denote significant effects, with α equal to 0.05, and levels varying according to probability ranges: *P* < 0.05 = *, *P* < 0.01 = **, *P* < 0.001 = ***.

### *Wolbachia* largely attenuates saliva transmission of DENV e ZIKV by field samples

To further understand the extent of *Wolbachia*-mediated refractoriness to ZIKV and DENV, we investigated the infective potential of saliva from orally-challenged Urca and Jurujuba samples. At 14 dpi, saliva samples were harvested and intrathoracically injected into groups of naïve Urca specimens (n=8) to evaluate the degree of which infective particles could be transmitted (Figure 6). Infected individual counts were assessed at 5 dpi, and their percent representation in each group was the metric used for comparisons between saliva samples. We also measured ZIKV and DENV titers in the heads of which saliva were harvested, and plotted the values at the top of each infected group. As previously observed in extracts following oral-infection, ZIKV and DENV titers were high in head samples from Urca, but virtually undetectable in those from Jurujuba (except for one sample with low titer). However, it is important to note that these titers reflect the background infection status, not necessarily translating to the saliva.

**Figure 6.**
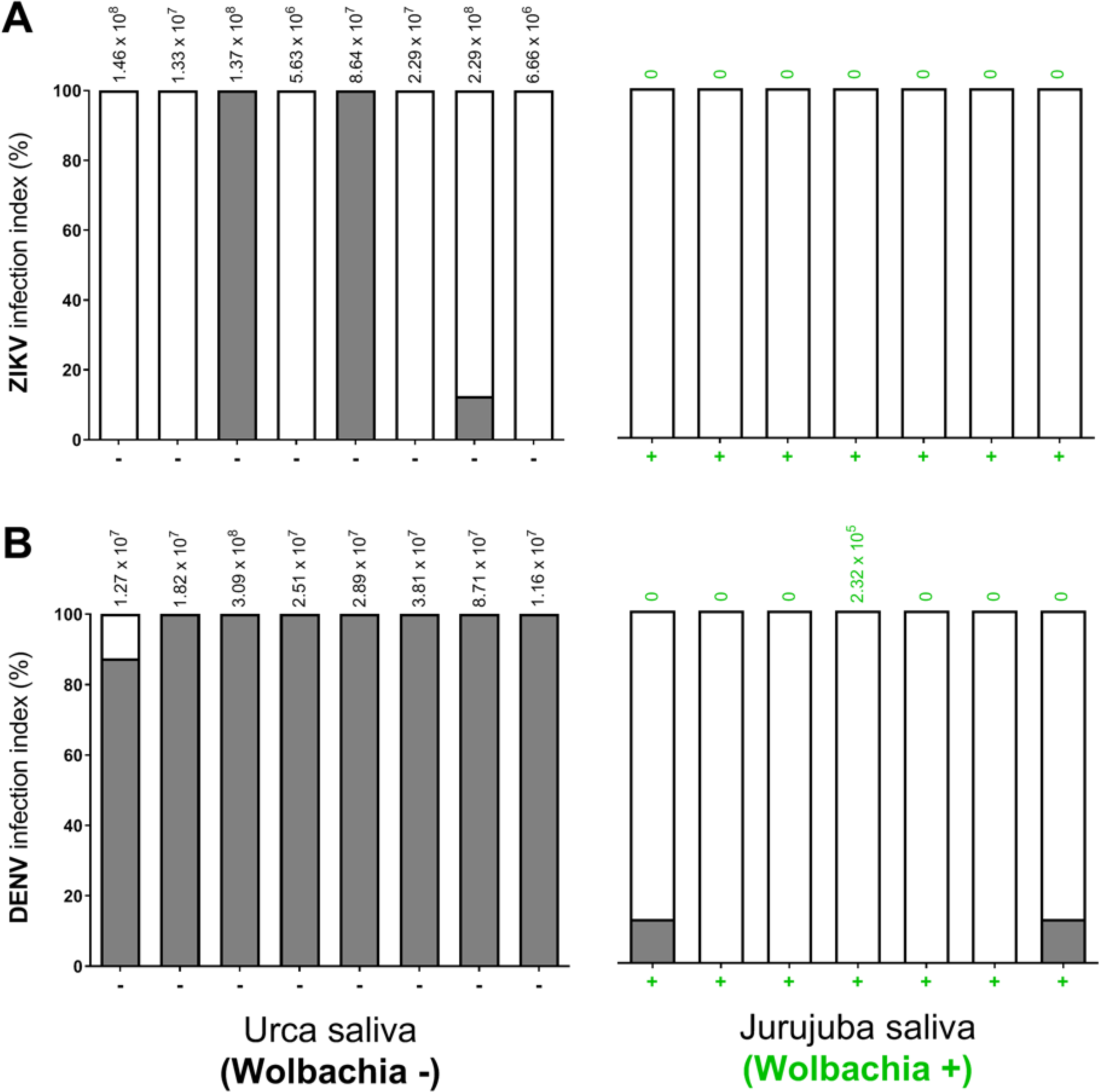
*Wolbachia*-harboring field samples impair saliva transmission of ZIKV and DENV. ZIKV and DENV transmission by the saliva is affected by the presence of *Wolbachia* in Jurujuba. Saliva harvested from *Wolbachia*^*−*^ Urca (grey) or *Wolbachia*^+^ Jurujuba samples (green), orally-challenged with ZIKV (A) or DENV (B), were intrathoracically injected into groups of naïve Urca individuals (n=8) and consequent infection was determined at 5 dpi. Groups are represented by stacked columns of infected (filled in grey) or non-infected individuals (clear), percent transformed. Viral titers of head extracts, of corresponding saliva samples, are shown at the top of each column.

Our results revealed that saliva samples from orally-infected Urca individuals were able to transmit ZIKV and DENV infectious particles. The transmission rates for ZIKV and DENV, however, seemed to differ. While 3 out of the 8 groups challenged with ZIKV-infected saliva had at least one individual positive for the virus (Figure 6A), all 8 groups challenged with DENV-infected saliva elicited this response (Figure 6B). This evidence suggests that despite being susceptible to both viruses, Urca population might be less competent to transmit ZIKV than DENV.

In contrast, saliva samples from orally-infected Jurujuba individuals (*Wolbachia*^+^) were usually unable to transmit ZIKV and DENV infectious particles. In fact, all 7 groups challenged with saliva from ZIKV-infected individuals did not elicit a single infection (Figure 6A), whilst only 2 out of 7 groups challenged with saliva from DENV-infected individuals were positive for the virus (a single infection in each group) (Figure 6B). These findings support the view that the *Wolbachia*-harboring Jurujuba population has an extremely low susceptibility to ZIKV and DENV infection, with reduced viral multiplication and dissemination within key organismal tissues, not being able to efficiently transmit the virus through saliva inoculation.

## Discussion

The global burden of dengue, Zika and chikungunya places *Ae. aegypti* at the top of the list encompassing medically relevant mosquito vectors (Bhatt et al., 2013; Kotsakiozi et al., 2017; Kraemer et al., 2015). Since human immunization is not an option to date, public health authorities focus their efforts on vector suppression campaigns using long-standing protocols with major constraints. Mechanical removal of breeding sites, for instance, are labor intensive and usually leave some hotspots untouched, in which dry quiescent eggs remain viable until more favorable conditions resume (Diniz et al., 2017; Farnesi et al., 2015; Oliva et al., 2018). Deployment of chemical pesticides have also proven inefficient, given the lack of precision and the surge of resistant variants (Maciel-de-Freitas et al., 2014; Macoris et al., 2018). To tackle some of these constraints and fulfil the urgent need for more efficient strategies, innovative solutions have emerged in recent years (Burt, 2014; Garcia et al., 2019; Harris et al., 2012; Hoffmann et al., 2011; Ryan et al., 2019; Schmidt et al., 2017; Tantowijoyo et al., 2020).

One promising solution lies on the field-release of lab-reared *Wolbachia*-infected individuals to gradually replace wild uninfected populations. The concept is fundamentally based on both the CI and PI bacterium-driven effects, arising from complex interactions with the mosquito host (Caragata et al., 2016). While the first favors bacterium inheritance towards fixation, the second confers refractoriness to several arboviruses. Should these effects translate to wild populations, then *Wolbachia* offers an unprecedent strategy, both natural and sustainable, to control mosquito-borne disease transmission.

Over the last decade, with the aid of the World Mosquito Program (formerly ‘Eliminate Dengue: Our Challenge’), field-release trials have been performed to assess whether *Wolbachia* can live up to expectations in diverse real-world scenarios (Garcia et al., 2019; Hoffmann et al., 2011; Nazni et al., 2019; Ryan et al., 2019; Tantowijoyo et al., 2020). Interestingly, trials have revealed several challenges for a successful *Wolbachia* invasion and long-term stability in natural populations. As highlighted by pilot studies in Northern Australia and Vietnam, choosing a bacterium strain associated with high fitness costs to the host, like the virulent *w*MelPop, may impair a population replacement strategy (Nguyen et al., 2015). Even though it could still be used in alternative strategies to transiently suppress *Ae. aegypti* populations and local transmission of arbovirus (Ferguson et al., 2015; Ritchie et al., 2015), its field application is not sustainable and presumes continuous release of large quantities of individuals (Rasić et al., 2014). In contrast, the utilization of strains associated with low-to-mild fitness costs, albeit with generally lower PI, allow fewer individuals to be released in the field in order to promote an efficient invasion. Fulfilling these criteria, the strain *w*Mel has been advocated as a standard choice for field release, collecting successful trials in Australia, Brazil and Indonesia (Garcia et al., 2019; Hoffmann et al., 2014, 2011; Indriani et al., 2020; O’Neill et al., 2019; Ryan et al., 2019; Tantowijoyo et al., 2020). Nonetheless, due to its beneficial attributes in warmer climates, keeping density levels high and stable, *w*AlbB has proven a second option and suitable alternative to *w*Mel, as shown by a recent trial held in Malaysia (Nazni et al., 2019). Thus, the investigation and field application of new strains has been encouraged to broaden the available options, and help building a *Wolbachia* toolset that suits diverse needs (Ant et al., 2018).

In addition to the *Wolbachia* strain, other determining factors impacting the invasion dynamics, hence the success of a field trial, include the genetic background of the host, the rear and release method *per se* (space-time release schedule, quantity and quality/fitness of released individuals) and the density of local *Ae. aegypti* populations (Garcia et al., 2019; Ryan et al., 2019). Of particular importance, the genetic background of the host not only establishes unique interactions with *Wolbachia* strains, but also harbors bacterium-independent fitness traits that may dictate the adaptation to wild environments. Corroborating to this view, background homogenization was key to revert a failed attempt to deploy *w*Mel in Tubiacanca, a small community of Rio de Janeiro (RJ), Southeastern Brazil (Garcia et al., 2019). Here, mimicking prior successful trials held in Australia (Hoffmann et al., 2014, 2011) was not enough to drive a sustainable invasion, and soon after adult release ceased *Wolbachia* infection indexes started to decay. Neither increasing the quantity nor the quality of released individuals (i.e. larger, longer-living specimens) could revert this outcome, suggesting that fitness-related nuances could be limiting *Wolbachia*’s spread. After a thorough investigation of the infected line, ruling out putative variations in traits affecting mating and reproduction success (Farnesi et al., 2019; Garcia et al., 2019; Gesto et al., 2018), or *Wolbachia*’s maternal transmission rate, it was revealed that its genetic footprint of insecticide resistance had been largely attenuated over laboratory adaptation, becoming particularly less fit to survive in insecticide-loaded habitats like those found in Southeastern Brazil (Garcia et al., 2019). With the re-introduction of resistant alleles, matching the frequency found in local populations, the reformed line was then able to switch the negative trend to a successful invasion in a second trial (Garcia et al., 2019). Thus, by narrowing disparities between lab-reared infected lines and wild populations, genetic background homogenization has been perceived as a good practice prior to current field release efforts.

In this study, we reported the successful introduction and long-term establishment of *Wolbachia* into *Ae. aegypti* populations from Jurujuba, a suburban community of Niterói (RJ). Sitting by the shores of Guanabara bay, Jurujuba represents an additional site for *Wolbachia* release trials in Rio de Janeiro (RJ) and surrounding areas, launched some years before in Tubiacanga (Garcia et al., 2019). Jurujuba is enclosed in a greener landscape with softly connected housing clusters (Figure 1), and Tubiacanga in a more organized and uniform housing display. Contrary to the latter, in which adult-releases were undertaken, Jurujuba held an alternative egg-release method, providing a chance to evaluate challenges and outcomes of each strategy, and trace future perspectives of trials in the country. The relatively small distance between both sites (19 km on a straight line over water) also proved valuable by allowing the field release of the same *Wolbachia*-infected line (i.e. *w*MelRio) (Garcia et al., 2019), disregarding the need for a whole genetic background swap. Instead, we carried out only minor quality control (e.g. genetic monitoring of insecticide resistance alleles) and re-introduction of local genetic variability in every few generations (e.g. addition of wild-caught males to the colony) to secure background homogeneity.

Following a rollout strategy, *Wolbachia*-containing eggs were deployed across all seven sectors of Jurujuba in over 73 weeks (or 58 weeks, if subtracted a 15-week inactive period between the release cessation in Ponto Final and start in Cascarejo). By monitoring the infection of field specimens, we revealed an overall invasion trend characterized by sustained high indexes at the end of the release period. However, a thorough observation of each sector uncovers distinct invasion profiles (Figure 2), with some featuring accentuated trends and reaching high infection indexes (>80%) in 5-17 weeks (e.g. Praia de Adão e Eva, Brasília, Peixe Galo and Ponto Final), and others showing less pronounced trends and reaching similar values in 24-37 weeks (e.g. Salinas and Cascarejo). An exception, not fitting into any of these categories, Várzea exhibited an erratic profile with a steep rise to high indexes during the release period, followed by mid-to-low index oscillation over 10 months before recovering and stabilizing at high indexes. The phenomena underlying distinct *Wolbachia* invasion dynamics could be linked to the density and spatial distribution of local *Ae. aegypti* populations and, ultimately, to human occupation (Hancock et al., 2016; Schmidt et al., 2017). Thus, it is reasonable to speculate that sectors featuring accentuated invasion trends have smaller populations of *Ae. aegypti*, as a result of better management of breeding sites or even fewer inhabitants. An opposing scenario might explain less pronounced, more resilient, invasion trends. Speculating on Várzea’s erratic profile, however, is challenging since none of the above causes seem reasonably suited to explain it assertively, leaving room to non-controlled events (e.g. indoors insecticide-spraying). Interestingly though, Várzea constitutes a stretch of houses surrounded by forest, and connecting three other sectors: Brasília, Cascarejo and Salinas (Figure 1). Thus, it is possible that migration from these adjacent sectors could have contributed to Várzea’s profile, first with non-infected individuals and then with infected ones, respectively dampening and recovering rates over the post-release phase.

Our data corroborate to a stable, self-sustaining, and long-term persistence of *Wolbachia* infection in *Ae. aegypti* from Jurujuba. We have shown that over the post-release phase, spanning mid-Juanuary 2017 to December 2019, *Wolbachia* infection of field specimens were sustained at near-fixation indexes with only minor fluctuation (80-100%) (Figure 3). Curiously, while experiencing and adapting to the natural habitat, *Wolbachia*’s association with infected hosts seems to have evolved to higher whole-body densities (Figure 4). When a comparison is drawn to colony-reared individuals, a significant increase in this parameter can be observed after only a few months (i.e. 2-4) in the field, becoming even more pronounced after 1 year. Considering that density levels has been positively correlated to maternal transmission rates (Ant et al., 2018; Ross et al., 2017), as well as to the strength of CI and PI (Lu et al., 2012; Martinez et al., 2014; Osborne et al., 2012; Terradas and McGraw, 2017; Walker et al., 2011), our findings suggest *Wolbachia*-host association affects have not been alleviated due to co-evolution in the field, and shall endure in the years to come.

Indeed, further supporting this view, our vector competence analysis suggests that PI is maintained in *Wolbachia*-positive Jurujuba samples collected slightly over 1-year into the post-release phase (Figure 5, Figure 6). In both oral and saliva challenging assays, Jurujuba samples were highly refractory to ZIKV or DENV, impairing not only the multiplication of viral particles but also its dissemination to organs playing key roles in transmission to humans (e.g. salivary glands). Since most samples from Jurujuba were *Wolbachia*-positive (∼ 88%) (Supplementary Figure 3) and those from Urca, a *Wolbachia*-free area, were highly susceptible to both viruses, we could endorse that the refractory effect was bacterium driven. Despite numerous studies of pathogen interference in *w*Mel-harboring lines, including some on a Brazilian background context (Dutra et al., 2016; Pereira et al., 2018), the data presented here account for the first evidence of ZIKV- and DENV-blocking in samples from Rio de Janeiro and surrounding areas, notably Jurujuba and Tubiacanga, which have been subjected to release trials in recent years. Most important, it adds an important validation to the undergoing control strategy, leaving to epidemiological analysis the last verdict.

Corroborating previous trials in Australia and Indonesia (O’Neill et al., 2019; Ryan et al., 2019; Tantowijoyo et al., 2020), the release of *Wolbachia*-infected eggs in Jurujuba proved an efficient method to introduce and disseminate the bacterium into Brazilian populations of *Ae. aegypti*. When compared to adult-driven methods, egg releases thrive on its simpler, more natural approach. Replacing a large, all-at-once, pulse of mosquitoes by a slow and gradual release from MRCs tends to alleviate undesirable social effects and increase community acceptance (Tantowijoyo et al., 2020). Interestingly, volunteers and community members can actively participate, with low-demanding basic training, in field deployment schedules, naturally enhancing engagement and the strategy’s successful outcome (O’Neill et al., 2019). Altogether, these beneficial features highly encourage the release of *Wolbachia*-infected eggs as part of control strategies in Brazil and other countries, particularly in those sites lacking proper infra-structure or financial support.

Ultimately, this work adds a new chapter on a successful story of *Wolbachia* field release in Southeastern Brazil. When associated with a local genetic background, and continually monitored for homogeneity, *w*Mel’s costs on fitness could be overridden by its efficient drive mechanism and spread into wild populations of Rio de Janeiro. Its long-term stability in the field, as shown by persistent high-infection indexes and pathogen interference, further reinforces the method’s sustainability and constitutes solid grounds to future epidemiological studies. Should we observe a significant impact on humans, then *Wolbachia*’s deployment shall gain momentum in public health initiatives and pave the way to cover larger areas in the country.

## Methods

### Mosquito lines and maintenance

To introduce *Wolbachia* into Brazilian *Ae. aegypti*, an Australian line infected with the *w*Mel strain (Walker et al., 2011) was backcrossed for 8 generations to a natural mosquito population of Rio de Janeiro, Brazil (Dutra et al., 2015). Following the genetic background swap, additional crosses and *knockdown resistance* (*kdr*) screening were undertaken to replicate natural insecticide resistance profiling and generate the line *w*MelRio. To assure a minimal variation in this profiling overtime, and sustain a homogeneous genetic background, *w*MelRio colony was refreshed with 10% wild males once in every five generations (Garcia et al., 2019).

To maintain *w*MelRio, immatures (ie. larval stages L1 to L4) were reared in dechlorinated water, at 28 °C, and fed Tetramin^®^ Flakes (Tetra GmbH, Herrenteich, Germany) until pupal formation. Following adult emersion, groups of 1,000 females and 800 males were sorted and kept in BugDorm^®^ cages (MegaView Science Co Ltd, Taiwan) at 25 °C, with 10% sucrose solution *ad libitum*. Every three days, females were fed human blood (from blood donation centers; see details under ‘ethical considerations’), through Hemotek^®^ artificial feeders (Hemotek Ltd, UK). Note that, to avoid arboviral contamination of our colony, all blood samples were formerly tested negative for dengue, Zika, chikungunya, mayaro and yellow fever by multiplex qPCR assays (Dutra et al., 2016; Pereira et al., 2018). Egg-laying was induced by placing dampened strips of filter paper (ie partially immersed in water-containing cups) inside the cages for 2-3 days, after which they were gradually dried at room temperature. Strips loaded with eggs (ie. ovistrips) were kept at room temperature until further use, either for colony maintenance or field release. Eggs older than 40 days were discarded due to a decay in overall quality (Farnesi et al., 2019).

### Egg releases

Mass-reared *w*Mel-infected Brazilian *Ae. aegypti, w*MelRio, were released as eggs in Jurujuba (22°56’00’’S, 43°07’00’’W), a lower-middle-class community in the city of Niterói (state of Rio de Janeiro, Brazil). Located by the shores of Guanabara bay, this community has grown from a typical fisherman settlement, with informal occupancy, to a total population of 2,797 residents in 890 houses. Jurujuba encompasses a total area of 2.53 km^2^, divided into seven smaller sectors: Ponto Final, Várzea, Brasília, Cascarejo, Praia de Adão e Eva, Peixe-Galo and Salinas.

*w*MelRio eggs were released in the field through special deployment devices, referred to as mosquito release containers (MRCs), which consisted of small white plastic buckets (19 cm height x 18 cm top diameter x 15.5 cm base diameter) with four small holes on the side wall, only a few centimeters away from the top lid. Each MRC was loaded with 1 L of water, 0.45g of Tetramin^®^ Tropical Tablets (ie. one and a half tablet) (Tetra GmbH, Herrenteich, Germany) and an ovistrip containing approximately 150 eggs. After 6 to 7 days, about 80% of the immatures were pupae, and after 11 to 12 days, most of the adults had already emerged and left the device from the wall holes. Every 15 days, MRCs were checked and reloaded so that another rearing and release cycle could take place. Release sites were spatially distributed as evenly as possible (Supplementary Figure 1), so as to maximize the spread of *Wolbachia*-harboring individuals and promote mating with their wild peers. The release strategy was optimized by splitting the sites into two groups, A and B, with alternate MRC loading schedules. Thus, while MRCs from group A were releasing adults, those from group B were being loaded with new ovistrips, water and food. In the following week, an opposite situation occurred, with MRCs from group B releasing adults. The release schedules, as well as the number of allocated MRCs, varied according to each Jurujuba’s sector (Supplementary Table 1).

### Ethical considerations

Regulatory approval for *Wolbachia* field release were obtained from the National Research Ethics Committee (CONEP, CAAE 02524513.0.1001.0008) and three government agencies: IBAMA (Ministry of Environment), Anvisa (Ministry of Health) and MAPA (Ministry of Agriculture, Livestock and Supply) to obtain the RET (Special Temporary Registry, 25351.392108/2013-96). Approval by CONEP implied a formal consent of at least 70 % of households, and that the World Mosquito Program would not change the usual practices performed at the community by the municipality vector control agents.

For the maintenance and mass-rearing of *Wolbachia*-infected *Ae. aegypti*, adult females were fed human blood from a donation center (Hospital Antonio Pedro, Rio de Janeiro State University), with supporting regulatory approval (CONEP, CAAE 59175616.2.0000.0008) We only used blood bags which would have been discarded by the donation center, mainly due to insufficient volume to meet their quality assurance policy. Samples had no information on donor’s identity, sex, age and any clinical condition, but were tested negative for several diseases, including Hepatitis B, Hepatitis C, Chagas disease, syphilis, HIV and HTLV, as part of the Brazilian Government routine screening.

For vector competence assays, human blood was obtained from Fundação Hemominas as part of a research agreement with Instituto René Rachou (Fiocruz Minas) (OF.GPO/CCO-Nr224/16).

### *Wolbachia* field monitoring and density level assessment

*Ae. aegypti* field population was monitored with BG-Sentinel traps (Biogents AG, Regensburg, Germany), spread across Jurujuba in a semi-homogeneous fashion (Supplementary Figure 2, Supplementary Table 2). These monitoring sites were chosen among suitable households who formally agreed with hosting of a trap, and had to be reallocated according to necessity (ie. household quits hosting the trap). Working traps were checked weekly by removing the catch bags (eg. small meshed envelopes placed inside the BG-Sentinels to collect trapped insects) and bringing them to the laboratory for species identification and *Wolbachia* screening. Catch bags were barcoded according to the trap ID and site, so as to create a pipeline for field samples.

*Ae. aegypti* positive samples were molecularly screened for *Wolbachia* by qPCR. Briefly, individual DNA was extracted by homogenizing head/thorax body parts in Squash Buffer (10 mM Tris-Cl, 1 mM EDTA, 25 mM NaCl, pH 8.2) supplemented with Proteinase K (200 ug/ml) and incubating at 56 °C for 5 min. Extraction ended by enzyme inactivation at 98 °C for 15 min. DNA amplifications were carried out with FastStart Essential DNA Probes Master (Roche), using specific primers and probes to *Wolbachia pipientis* WD0513 and *Ae. aegypti rps17* markers (Supplementary Table 3). Thermocycling conditions were set on a LightCycler^®^ 96 Instrument (Roche), as follows: 95 °C for 10 min (initial denaturation), and 40 cycles of 95 °C for 15 s and 60 °C for 30 s. *Wolbachia* density levels were defined as a relative index corresponding the ratio of WD0513 and RPS17 amplified products.

### DENV and ZIKV isolation and replication in mosquito cells

ZIKV was kindly provided by Instituto Aggeu Magalhães (IAM, Fiocruz) through viral isolation of a symptomatic patient sample from Recife (PE, Brazil) in 2015 (ZikV/ H.sapiens/ Brazil/ BRPE243/ 2015). DENV was sourced by our group at Instituto René Rachou (IRR, Fiocruz), following a viral isolate from a patient sample diagnosed with Dengue type 1 in Contagem (MG, Brazil), also in 2015 (Den1/ H.sapiens/ Brazil/ BRMV09/ 2015). Both ZIKV and DENV samples were accompanied by patients’ written consent (CONEP, reference number 862.912), being further catalogued into the national database of genetic patrimony and associated knowledge (SISGEN, access number AA1D462).

*In vitro* culture of viral particles were done as previously described (Dutra et al., 2016). Briefly, ZIKV and DENV were replicated in *Aedes albopictus* C6/36 cells, grown at 28 °C in Leibovitz L-15 medium (ThermoFisher) supplemented with 10% fetal bovine serum (FBS) (ThermoFisher). After seven days, supernatants were harvested and virus titers were assessed, first by Reverse Transcription (RT)-qPCR, and later by plaque assay with VERO cells grown under 37 °C, 5% carbon dioxide, in Dulbecco’s Modified Eagle Medium (DMEM) (ThermoFisher) supplemented with 3% Carboxymethylcellulose (Synth) and 2% FBS.

### Vector competence assays

To perform vector competence assays with field samples of *Ae. aegypti*, ovitraps were mounted in both Ponto Final (Jurujuba) and Urca, a *Wolbachia*-free area in Rio de Janeiro. Ovitraps were collected from the field over 13 weeks, from April to June 2017, which corresponds to the time-frame between 14 and 16 months along the post-release phase in Ponto Final. Once in the insectary, eggs samples were reared to the adult stage in a controlled insectary environment (refer to ‘mosquito lines and maintenance’ for details).

For virus challenging assays through oral-feeding, young females (4-6 days old) were starved for 20 to 24 h, and subsequently offered culture supernatant containing ZIKV or DENV mixed with human red blood cells (2:1 ratio), using an artificial membrane feeding system (Dutra et al., 2016). It is important to mention that, as for the colony maintenance protocol, blood samples used here were also submitted to quality control prior to its use in the assays, mainly due to putative arbovirus contaminations which could affect the experimental outputs. Likewise, all samples were tested negative for dengue, Zika, chikungunya, mayaro and yellow fever by multiplex qPCR assays (Dutra et al., 2016; Pereira et al., 2018). Oral-infections were performed twice for each virus. ZIKV was offered first from fresh (virus titer of 4.8 × 10^8^ PFU/mL) and second from frozen culture supernatant (virus titer of 7.6 × 10^6^ PFU/mL). In contrast, DENV was offered from fresh supernatants only (virus titers of 2 × 10^6^ PFU/mL and 6.5 × 10^7^ PFU/mL), since frozen versions failed to infect. Specimens were allowed to feed for one hour, after which engorged females were selected and maintained with 10% sucrose solution *ad libitum*, during the whole extrinsic incubation period. At 14 days post-infection (dpi), viral loads were assessed in heads/thorax extracts by RT-qPCR (refer to ‘Viral diagnosis’ for more details).

For saliva-mediated virus challenging assays, ZIKV and DENV pre-exposed females (14 dpi) from Jurujuba (*Wolbachia* +) and Urca (*Wolbachia* -) were starved for about 16 h (overnight) before being knocked down and kept at 4°C for wings and legs removal. Salivation was induced by introducing a 10 µL sterile filter tip, pre-loaded with 5 µl of 50% fetal bovine serum (FBS) and 30% sucrose, into the mosquito proboscis for 30 min. Saliva samples were individually titrated, and 276 nL was intrathoracically injected into young naive females (Urca) using a Nanoject II hand held injector (Drummond), as previously described (Dutra et al., 2016; Pereira et al., 2018). Each saliva sample was used to inoculate 8-14 naïve *Wolbachia*-free *Ae. aegypti* specimens, of which 8 were screened for infective particles. ZIKV and DENV were quantified by RT-qPCR at 5 dpi and 7 dpi, respectively (refer to ‘Viral diagnosis’ for more details).

### Viral diagnosis

To identify ZIKV and DENV particles in individual samples, whole specimens were processed into head/thorax homogenates for RNA extraction with the High Pure Viral Nucleic Acid Kit (Roche), according to manufacturer’s instructions (Moreira et al., 2009). Saliva RNA was extracted with the same reagents, though using half of the volumes advised. RNA samples were diluted in nuclease-free water to a concentration of 50 ng/μL. ZIKV, DENV and *Wolbachia* levels, in vector competence assays, were quantified by RT-qPCR using TaqMan Fast Virus 1-Step Master Mix (ThermoFisher) and specific primers and probes (Supplementary Table 3). Reactions were run on a LightCycler^®^ 96 Instrument (Roche), using the following thermocycling conditions: 50°C for 5 min (initial RT step), 95°C for 20 s (RT inactivation / DNA initial denaturation), and then 40 cycles of 95°C for 3 s and 60°C for 30 s. Each RNA sample was used in two reactions, one with ZIKV, DENV or *Wolbachia* primers, and another with *Ae. aegypti rps17* endogenous control (Moreira et al., 2009). Absolute quantification was achieved by comparing amplification profiles with standard curves generated by serial dilutions of their respective gene products, amplified from a cloned sequence in pGEM^®^-T Easy vector (Promega). Negative control samples (no virus DNA) served as reference to fix a minimum threshold for positive ones. ZIKV and DENV loads were defined as their copy number per sample (head/thorax or saliva), while *Wolbachia* loads were relative quantifications to the *rps17* marker.

### Statistical Analyses

Graphs and statistical analyzes were performed in Graphpad Prism 8 (Graphpad Software, Inc). Kruskal-Wallis test followed by Dunn’s post-hoc multiple comparisons were used to analyze *Wolbachia* density data from field-collected and colony samples. ZIKV and DENV loads in head/thorax extracts, from both oral and saliva-challenging samples, were compared using the Mann-Whitney U test. For all statistical inferences, α was set to 0.05.

## Acknowledgements

We would like to thank all members of the Mosquitos Vetores research group, for critical and helpful reviews, as well as past and present team members of the World Mosquito Program, whose dedication was key to the positive outcome of this study. Special thanks to Flavia Teixeira, for ethical and regulatory compliance, Roberto Peres and Catia Cabral, for field deployment and mass-rearing supervision and Simon Kutcher (WMP Global) for overall technical inputs, during early days of our implementation. We are also grateful for the field assistance provided by public agents of the Health Municipality of Niterói, and for the incredible support by Jurujuba community members.

## Figures and Tables

**Supplementary Figure 1.**
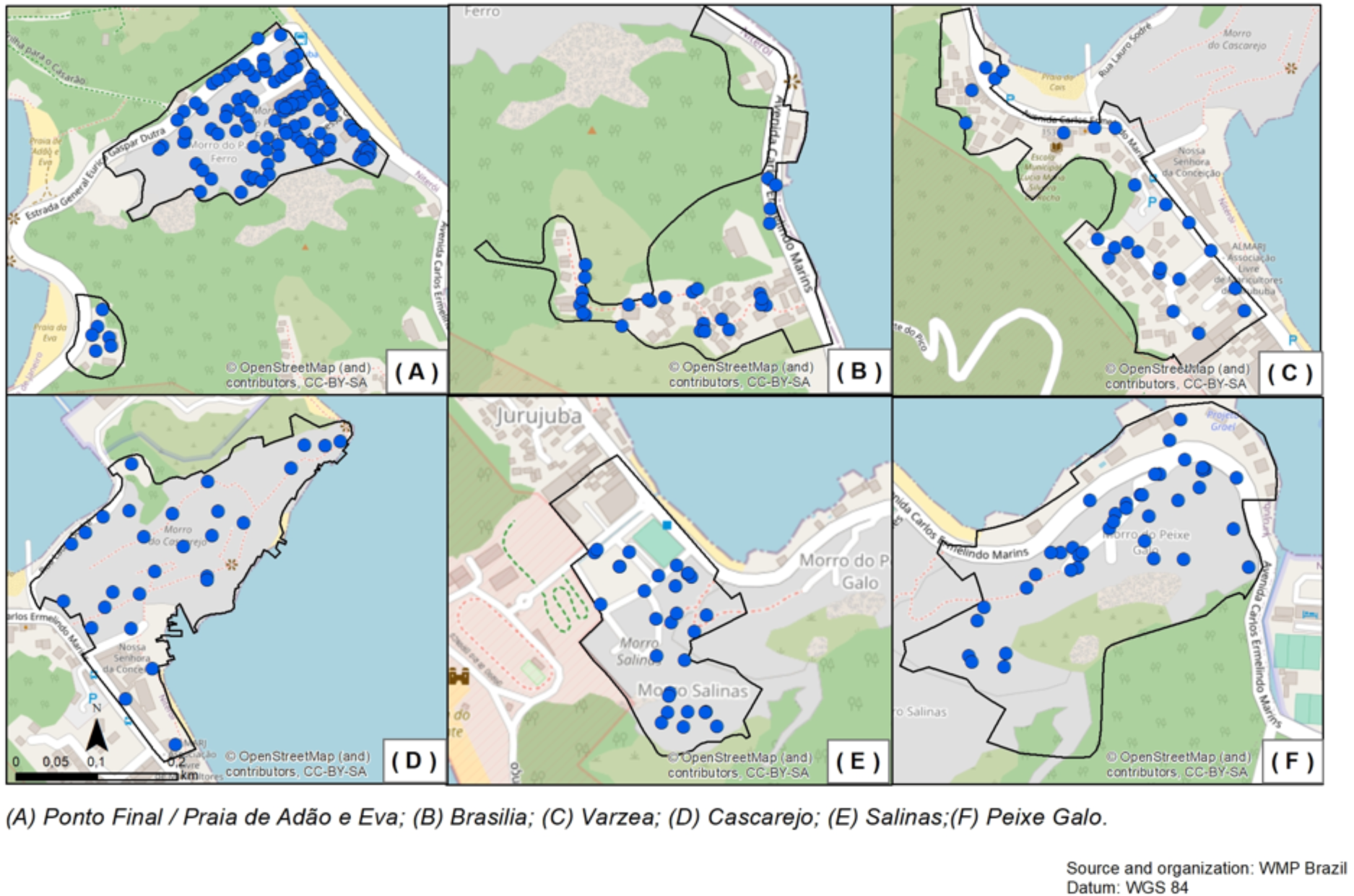
Map of egg-release sites. Spatial distribution of Mosquito Release Containers (MRCs) (blue circles) across Jurujuba’s sectors. At 15-days intervals, each MRC was loaded with a fresh batch of *w*Mel-infected eggs.

**Supplementary Figure 2.**
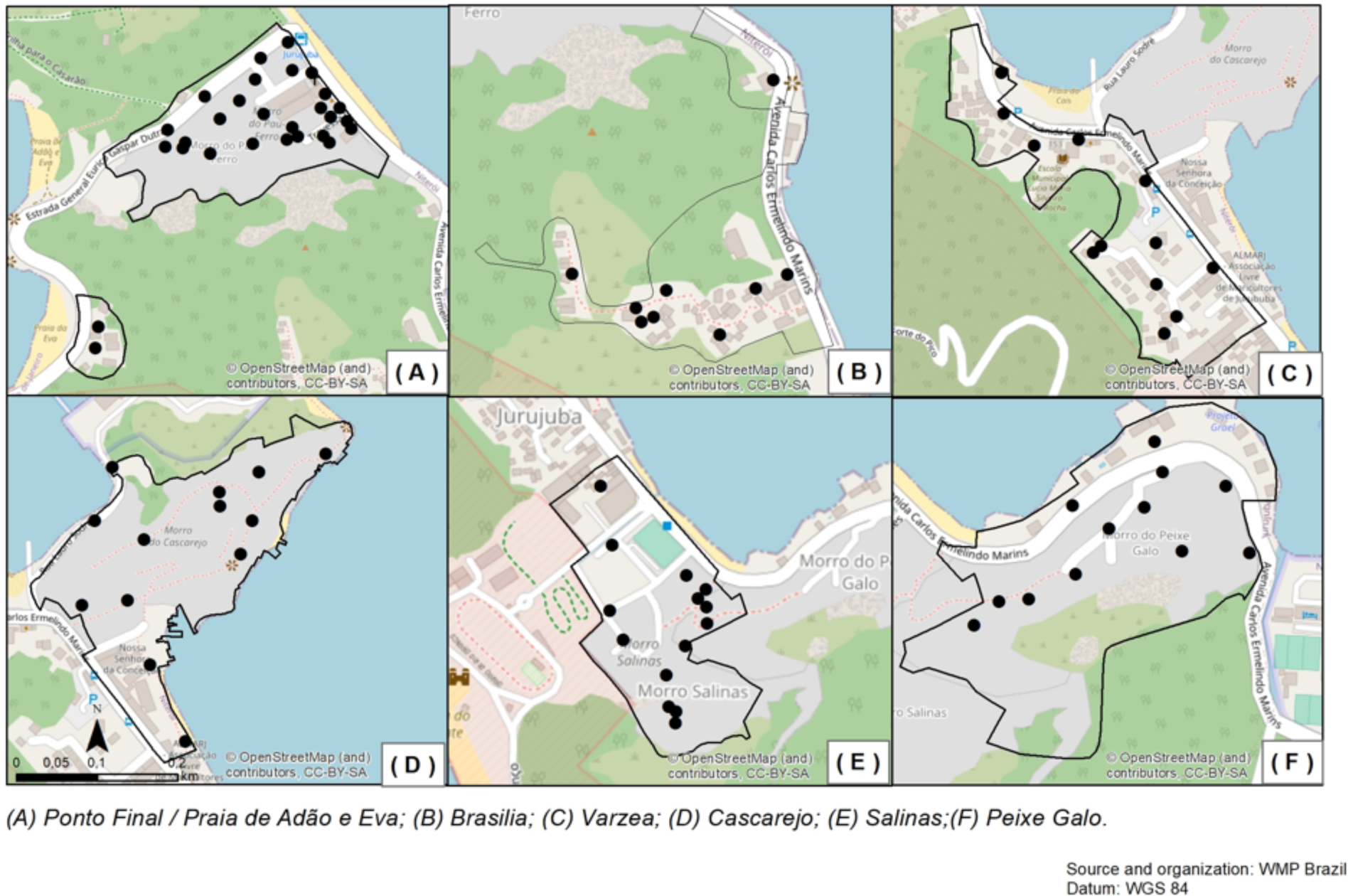
Map of monitoring sites. Spatial distribution of BG-sentinel traps (black circles) across Jurujuba’s sectors. Traps were monitored weekly to assess *Wolbachia* frequency in field specimens.

**Supplementary Figure 3.**
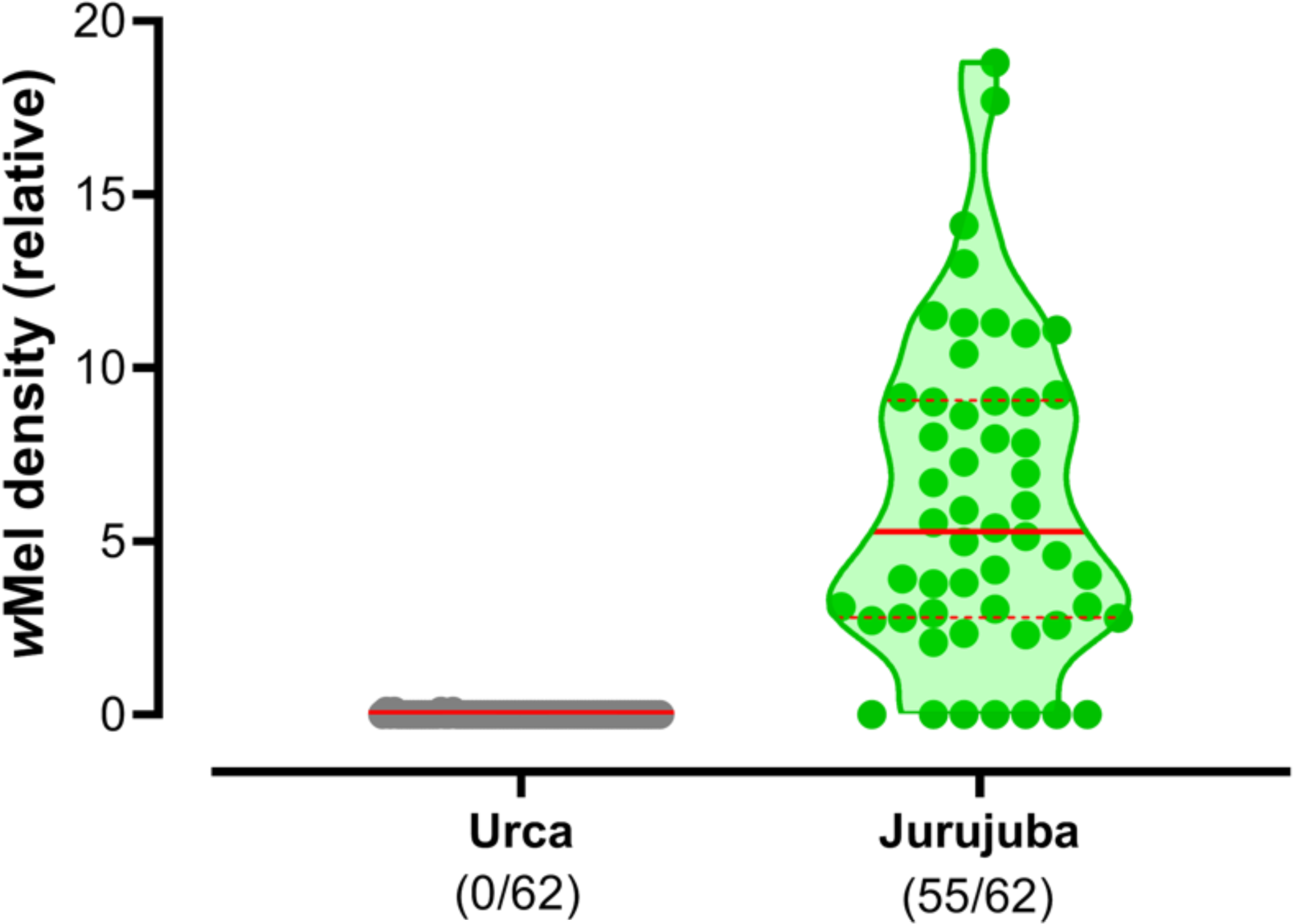
*Wolbachia* prevalence and density in virus-challenged samples. *Wolbachia* frequency and tissue density was assessed in Urca (grey) and Jurujuba samples (green) orally-challenged with ZIKV and DENV. Tissue density is a relative quantity, expressed as the ratio between *Wolbachia*-specific *WSP* and endogenous *RPS* genetic markers. Violin plots depict the distribution of density data from Urca and Jurujuba samples, with the proportion of infected individuals shown as fractions underneath. Medians are shown in solid and quartiles in dashed red lines.

**Supplementary Table 1.** Egg release schedules in Jurujuba. Along the *Wolbachia* field deployment period, Mosquito Release Containers (MRCs) were set up across Jurujuba’s territory in a variable fashion for each of its sectors. The release schedules and the number of MRCs allocated per sector are listed.

| Jurujuba's sector | Number of MRCs |  |  | Release Schedule |  |  |
| --- | --- | --- | --- | --- | --- | --- |
|  | Group A | Group B | Total | Start | End | Length (weeks) |
| Ponto Final | 57 | 58 | 115 | 25/08/2015 | 16/02/2016 | 25 |
| Cascarejo | 28 | 28 | 56 | 07/06/2016 | 10/01/2017 | 31 |
| Brasília | 14 | 14 | 28 | 13/09/2016 | 29/11/2016 | 11 |
| Salinas | 30 | 30 | 60 | 13/09/2016 | 03/01/2017 | 16 |
| Varzea | 12 | 12 | 24 | 13/09/2016 | 08/11/2016 | 8 |
| Peixe Galo | 21 | 21 | 42 | 20/09/2016 | 03/01/2017 | 15 |
| Praia de Adão e Eva | 6 | - | 6 | 20/09/2016 | 11/01/2017 | 16 |

**Supplementary Table 2.** BG-Sentinel traps allocation in Jurujuba. Number of BG-Sentinel traps allocated per Jurujuba sector, since the beginning of *Wolbachia* field monitoring until recent days. Traps were widely used and reached their maximum numbers at deployment periods, when monitoring with the best possible spatial resolution was critical, and then partially demobilized into the post-release phase following a successful *Wolbachia* invasion (ie. stable high-frequency infection indexes). Revision of schedules and trap numbers was necessary to allow a viable long-term monitoring activity across Jurujuba’s territory.

| Jurujuba's sector | Number of BG-traps |  | Monitoring schedule |  |
| --- | --- | --- | --- | --- |
|  | Max. | Min. | Start | End |
| Ponto Final | 26 | 3 | 08/09/2015 | 30/12/2019 |
| Cascarejo | 13 | 2 | 21/06/2016 | 30/12/2019 |
| Brasília | 9 | 2 | 27/09/2016 | 30/12/2019 |
| Salinas | 14 | 4 | 27/09/2016 | 11/12/2018 |
| Varzea | 11 | 2 | 27/09/2016 | 30/12/2019 |
| Peixe Galo | 12 | 1 | 27/09/2016 | 11/12/2018 |
| Praia de Adão e Eva | 2 | 2 | 06/10/2016 | 14/02/2017 |

**Supplementary Table 3.**
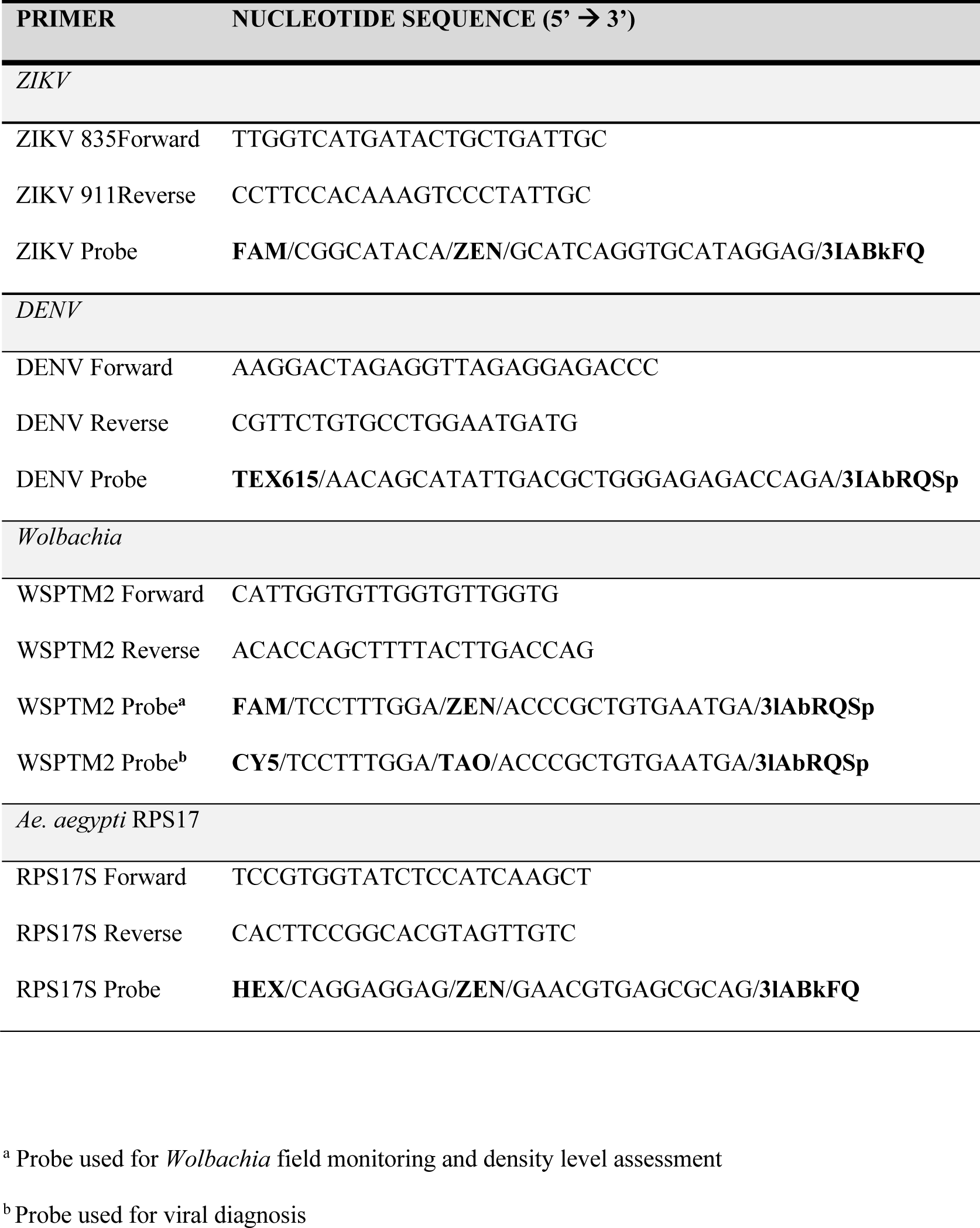
qPCR primes and probes. List of primers and probes for the molecular detection and quantification of ZIKV, DENV and *Wolbachia*.

